# TRAF6 and TAK1 contribute to SAMHD1-mediated negative regulation of NF-κB signaling

**DOI:** 10.1101/2020.11.07.368704

**Authors:** Constanza E. Espada, Corine St. Gelais, Serena Bonifati, Victoria V. Maksimova, Michael P. Cahill, Sun Hee Kim, Li Wu

## Abstract

Sterile alpha motif and HD-domain–containing protein 1 (SAMHD1) restricts HIV-1 replication by limiting the intracellular dNTP pool. SAMHD1 also suppresses the activation of NF-κB in response to viral infections and inflammatory stimuli. However, the mechanisms by which SAMHD1 negatively regulates this pathway remain unclear. Here we show that SAMHD1-mediated suppression of NF-κB activation is modulated by two key mediators of NF-κB signaling, tumor necrosis factor (TNF) receptor-associated factor 6 (TRAF6) and transforming growth factor-β-activated kinase-1 (TAK1). We compared NF-κB activation stimulated by interleukin (IL)-1β in monocytic THP-1 control and SAMHD1 knockout (KO) cells with and without partial TRAF6 knockdown (KD), or in cells treated with TAK1 inhibitors. Relative to control cells, IL-1β-treated SAMHD1 KO cells showed increased phosphorylation of the inhibitor of NF-κB (IκBα), an indication of pathway activation, and elevated levels of *TNF-α* mRNA. Moreover, SAMHD1 KO combined with TRAF6 KD or pharmacological TAK1 inhibition reduced IκBα phosphorylation and *TNF-α* mRNA to the level of control cells. SAMHD1 KO cells infected with single-cycle HIV-1 showed elevated infection and *TNF-α* mRNA levels compared to control cells, and the effects were significantly reduced by TRAF6 KD or TAK1 inhibition. We further demonstrated that overexpressed SAMHD1 inhibited TRAF6-stimulated NF-κB reporter activity in HEK293T cells in a dose-dependent manner. SAMHD1 contains a nuclear localization signal (NLS), but an NLS-defective SAMHD1 exhibited a suppressive effect similar to the wild-type protein. Our data suggest that the TRAF6-TAK1 axis contributes to SAMHD1-mediated suppression of NF-κB activation and HIV-1 infection.

**Importance:** Cells respond to pathogen infection by activating a complex innate immune signaling pathway, which culminates in the activation of transcription factors and secretion of a family of functionally and genetically related cytokines. However, excessive immune activation may cause tissue damage and detrimental effects on the host. Therefore, in order to maintain host homeostasis, the innate immune response is tightly regulated during viral infection. We have reported SAMHD1 as a novel negative regulator of the innate immune response. Here, we provide new insights into SAMHD1-mediated negative regulation of the NF-κB pathway at the TRAF6-TAK1 checkpoint. We show that SAMHD1 inhibits TAK1 activation and TRAF6 signaling in response to proinflammatory stimuli. Interestingly, TRAF6 knockdown in SAMHD1-deficient cells significantly inhibited HIV-1 infection and activation of NF-κB induced by virus infection. Our research reveals a new negative regulatory mechanism by which SAMHD1 participates in the maintenance of cellular homeostasis during HIV-1 infection and inflammation.

## Introduction

Sterile alpha motif and HD domain-containing protein 1 (SAMHD1) is a deoxynucleoside triphosphate triphosphohydrolase (dNTPase) that restricts human immunodeficiency virus type 1 (HIV-1) infection in non-dividing cells of myeloid lineage (1, 2) and resting CD4+ T cells (3, 4). Through its ability to hydrolyze deoxynucleoside triphosphates (dNTPs) (5, 6), SAMHD1 limits the amount of substrate available for replication of retroviruses and some DNA viruses (7-12). Germline loss-of-function mutations in the *SAMHD1* gene cause Aicardi-Goutières Syndrome (13, 14), which is manifested by increased production of type I interferon (IFN-I) (15). Furthermore, SAMHD1-deficient primary peripheral blood mononuclear cells isolated from patients with Aicardi-Goutières Syndrome are also highly permissive to HIV-1 replication (16). SAMHD1 is characterized as a predominantly nuclear protein, although it has been visualized in the cytoplasm in primary resting CD4+ T cells and macrophages (4).

We recently reported that SAMHD1 regulates the innate immune response through the suppression of nuclear factor-κB (NF-κB) activity induced by viral infections and inflammatory stimuli (17). We have shown that SAMHD1 deficiency in a THP-1 cell line increases NF-κB activation in response to known stimuli of the pathway (17), including Sendai virus (18) and the bacterial product and toll-like receptor 4 (TLR4) agonist lipopolysaccharide (LPS) (19). In addition, SAMHD1-mediated inhibition of NF-κB activation is dependent on its dNTPase activity, but not on its nuclear localization (20). NF-κB signaling is critical for the coordination of inflammatory processes (21), but the exact mechanism by which SAMHD1 inhibits the NF-κB pathway remains unknown (22).

NF-κB transcription factors remain sequestered in the cytoplasm by members of the inhibitory IκB family, and their activities are inducible through the canonical or non-canonical pathways (21). Activation of the canonical pathway is mediated by proinflammatory cytokines, including those of the interleukin (IL)-1 and tumor necrosis factor (TNF) families, and pathogen-associated molecular patterns, leading to the phosphorylation of the NF-kB inhibitor IκBα by the IκB kinase (IKK) complex. Subsequent ubiquitin-mediated proteasomal degradation of IκBα allows for nuclear translocation of the p50/RelA (p65) dimer (23). The non-canonical pathway differs by responding to a more selective set of stimuli and is dependent on the processing of the p100 precursor, mediated by NF-κB-inducing kinase (24).

Key upstream signaling events that result in the activation of IKK involve TNF receptor-associated factor 6 (TRAF6) and transforming growth factor-β-activated kinase-1 (TAK1, also known as MAP3K7). TRAF6 is a RING domain ubiquitin ligase that catalyzes Lys-63 (K63)-linked polyubiquitination of TAK1 at Lys158. TAB2 and TAB3 proteins bind to K63-linked polyubiquitin chains of TRAF6, resulting in the activation of TAK1 through auto-phosphorylation (25, 26). TAK1 is a mitogen-activated protein kinase kinase kinase (MAP3K) whose activity is induced by an array of proinflammatory stimuli, including IL-1, TNF-α and LPS. TAK1-mediated signaling activates NF-κB and the activator protein-1 transcription factors, which regulate many cellular processes, including cell proliferation and differentiation, survival, and innate and adaptive immune responses (27). Acting together in a signaling complex, TRAF6 and TAK1 may also direct the activation of mitogen-activated protein kinase (MAPK), one of several pathways that branch from TRAF6, independently of the IKK-NF-κB pathway. Therefore, the TRAF6-TAK1 axis serves as an important bridge between receptor signaling and NF-κB activation (28).

In this study, we observed a link between SAMHD1-mediated suppression of NF-κB activation and signaling events at the TRAF6-TAK1 axis. We show that endogenous SAMHD1 inhibits TAK1 phosphorylation and activation. Both wild-type (WT) SAMHD1 and its NLS-defective mutant inhibits TRAF6-mediated activation of NF-kB signaling, suggesting that SAMHD1 regulates NF-κB signal transduction in the cytoplasm. Furthermore, we demonstrate that SAMHD1 suppression of NF-kB activation during HIV-1 infection involves the TRAF6-TAK1 axis. The activation and suppression of innate immune responses is a dynamic balance, which is critical for defense against pathogens, prevention of autoimmunity and toxicity. Exploring how SAMHD1 contributes to this balance may provide insights toward the development of new treatment strategies to clear viral infections or control inflammatory diseases.

## Results

### Endogenous SAMHD1 inhibits TAK1 activation

Our previous gene expression analysis by Affymetrix microarray in THP-1 control and SAMHD1-deficient cell lines showed TAK1 is one of several host genes that has altered mRNA expression levels (29). Further analysis of the microarray data by Ingenuity Pathway Analysis (IPA) predicted that TAK1 is activated in SAMHD1-silenced THP-1 cells compared to control cells (Table 1). To validate our microarray data and IPA, the effect of SAMHD1 expression in THP-1 cells on the level of total and phospho-TAK1 (p-TAK1) at threonine 187 (T187) was examined by immunoblotting following IL-1β stimulation (Fig. 1A). We observed that total TAK1 protein levels were slightly decreased in SAMHD1 KO cells, which is expected based on the microarray data. Unstimulated SAMHD1 KO cells had 10-fold elevated p-TAK1 levels compared to control cells. Interestingly, p-TAK1 levels were 16-26-fold higher in IL-1β-stimulated SAMHD1 KO cells compared to unstimulated control cells. Moreover, (5Z)-7-Oxozeaenol (5Z), a selective and irreversible inhibitor of TAK1 (30), markedly decreased IL-1β-induced p-TAK1 (Fig. 1B) and *TNF-α* mRNA levels (Fig. 1C) in the THP-1 cell line, eliminating the enhancement of TAK1 activation observed with SAMHD1 deficiency. To confirm our results, we employed Takinib, a compound with higher selectivity for TAK1 (31). Reflecting the phenotype observed in our experiments with 5Z, the enhanced *TNF-α* mRNA levels in IL-1β-stimulated SAMHD1 KO cells were significantly reduced by pre-treating the cells with Takinib (Fig. 1D). Of note, neither 5Z nor Takinib treatment affected cell viability, assessed by the MTS cell proliferation assay (32) (data not shown). Altogether, these data suggest that endogenous SAMHD1 suppresses activation of the NF-κB canonical pathway by modulating TAK1 activity.

**Table 1.** Differential gene expression in SAMHD1 knockout (KO) THP-1 cells compared to control cells.

| Symbol | Entrez gene name | mRNA fold change | Expected activation<sup>a</sup> |
| --- | --- | --- | --- |
| KRAS | KRAS proto-oncogene, GTPase | -5.42 | Up |
| HDAC2 | Histone deacetylase 2 | -3.90 | Down |
| TAB2 | TGF-beta activated kinase 1/MAP3K7 binding protein 2 | -3.37 | Up |
| MAP3K7<sup>b</sup> | Mitogen-activated protein kinase kinase kinase 7 | -3.25 | Up |
| IGF2R | Insulin like growth factor 2 receptor | -2.95 | Up |
| PIK3R1 | Phosphoinositide-3-kinase regulatory subunit 1 | 2.23 | Up |
| PRKACB | Protein kinase cAMP-activated catalytic subunit beta | 2.12 | Up |
| PIK3C3 | Phosphatidylinositol 3-kinase catalytic subunit type 3 | 2.23 | Up |
| FCER1G | Fc fragment of IgE receptor Ig | 2.51 | Up |
| IL18 | Interleukin 18 | 3.49 | Up |
| CD40 | CD40 molecule | 3.63 | Up |
| TLR6 | Toll like receptor 6 | 3.68 | Up |
| AKT3 | AKT serine/threonine kinase 3 | 10.71 | Up |

<sup>a</sup> The expected activation of genes whose mRNA expression levels were significantly altered in THP-1 SAMHD1 KO cells relative to control cells (fold change > 2, $P \leq 1 \times 10^{-4}$ ) was predicted through Ingenuity Pathway Analysis (IPA) of the microarray data. RNA extracted from triplicate THP-1 control and SAMHD1 KO cells was analyzed by Affymetrix Clariom D Microarray (29).
<sup>b</sup> MAP3K7 is also known as transforming growth factor-β-activated kinase-1 (TAK1).

**Fig. 1.**
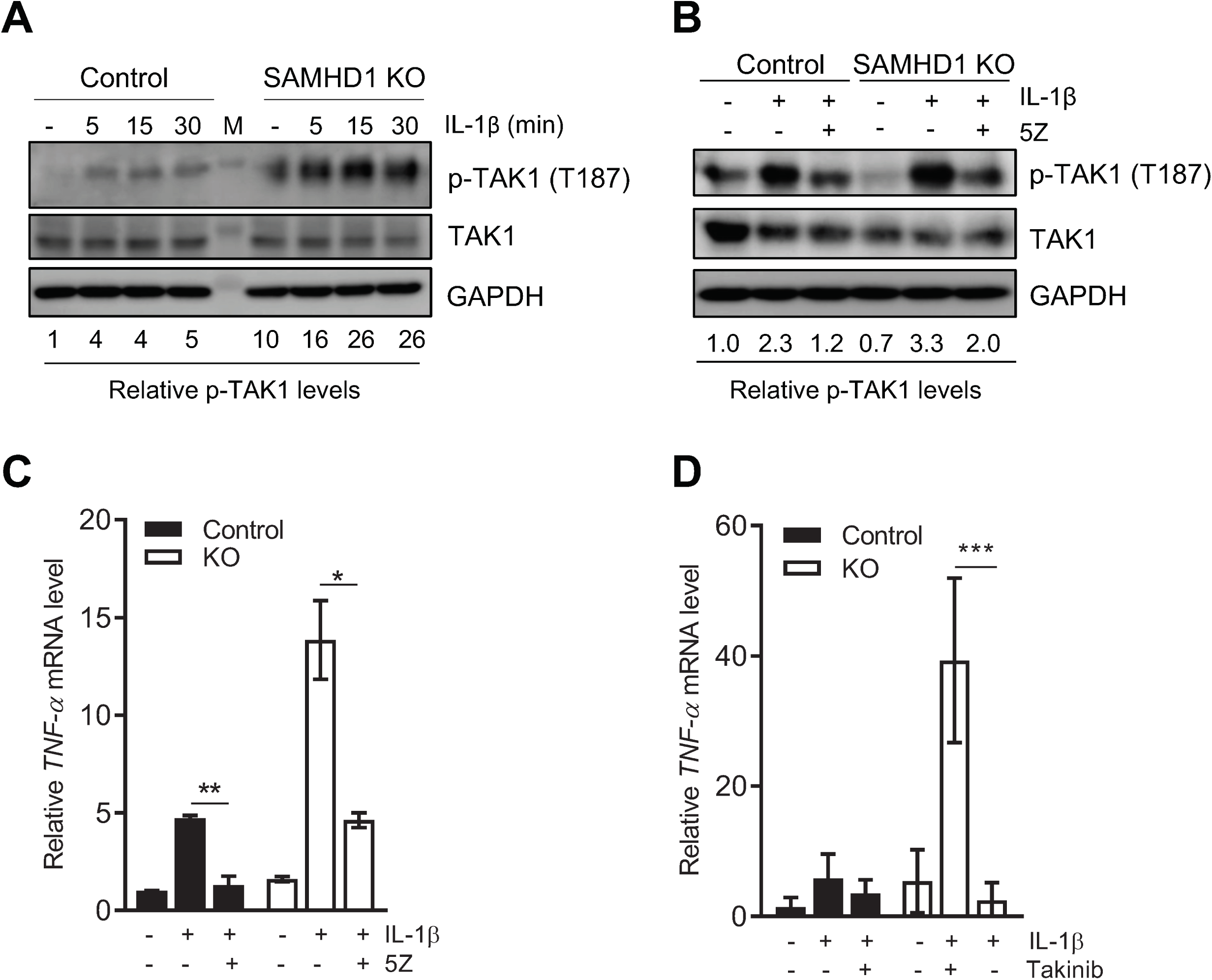
Endogenous SAMHD1 inhibits TAK1 activation. **(A)** THP-1 control and SAMHD1 KO cells were seeded in low glucose (5.5 mM) media for 48 hr. Cells were mock-treated or treated with IL-1β, harvested at 5, 15, and 30 min, and then lysed for immunoblotting. GAPDH was used as a loading control. Relative p-TAK1 proteins levels were calculated by densitometry analysis with normalization to total TAK1 levels and GAPDH. Mock-treated cells were set as 1. (**B**) THP-1 control and KO cells were grown as in (A). Cells were then cultured in 5Z (1 µM) or DMSO for 30 min. After 5Z removal, IL-1β was added to cells for 5 min prior to collection and analysis by immunoblotting, or for 2 hr for RT-qPCR detection of *TNF-α* mRNA in (C). Relative p-TAK1 protein levels were calculated by densitometry analysis. The p-TAK1 signal was normalized to total TAK1 protein and GAPDH. Untreated control cells (without inhibitor or IL-1β stimulation) were set as 1. (A-B) The immunoblots were representative data of three independent experiments. **(C)** Measurement of mRNA levels was performed from samples in the same experiment described in (B). *TNF-α* mRNA was normalized to spliced GAPDH. Data represent duplicate samples and error bars depict standard deviation (SD). Statistical significance was calculated by unpaired *t* test. **(D)** Cells were treated with Takinib (10 µM) for 2 hr prior to IL-1β stimulation. TNF-α mRNA was quantified by RT-qPCR as in (C). Data represents triplicate samples and error bars depict standard error of the mean (SEM). Statistical significance was calculated by unpaired *t* test. \**P* ≤ 0.05, \*\**P* ≤ 0.01, \*\*\**P* ≤ 0.001.

### Cytoplasmic SAMHD1 inhibits TRAF6-mediated activation of NF-kB signaling

TAK1 activation by IL-1β or stimulation of host pathogen recognition receptors (PRRs) depends on the recruitment of the upstream factor TRAF6, which catalyzes the formation of K63-linked polyubiquitin chains (33). Using a NLS-defective SAMHD1 mutant (mNLS), we have demonstrated that SAMHD1 inhibits IL-1β-induced NF-κB activation independently of its NLS (20). To determine if SAMHD1 suppress NF-κB activation initiated through TRAF6, we used an NF-κB-luciferase reporter plasmid system. Both WT SAMHD1 and mNLS inhibited NF-κB activation in a dose-dependent manner (Fig. 2A). Quantification of *luciferase (luc)* mRNA confirmed that SAMHD1-mediated suppression of NF-κB activation occurs prior to or at the level of mRNA transcription and that the nuclear localization of SAMHD1 is not required for this effect (Fig. 2B). To rule out general transcription inhibition by overexpressed SAMHD1, a plasmid expressing GFP under a cytomegalovirus promoter was co-transfected along with SAMHD1 and NF-kB luciferase reporter system. Co-expression of SAMHD1 did not inhibit the percentage of GFP-positive cells (Fig. 2C) or the level of GFP expression in transfected HEK293T cells (Fig. 2D), suggesting that SAMHD1-mediated inhibition of NF-kB activation is not due to a general suppression effect on cellular transcription. Our bioinformatic analysis revealed that the HD domain of SAMHD1 has a putative TRAF6 consensus-binding motif (^275^PXEXXXE^281^) (34) (Fig. 2E). To examine whether the potential binding of SAMHD1 and TRAF6 could affect TRAF6-mediated NF-κB activation, we generated a mutant SAMHD1 (P275A) with the altered sequence of the motif. However, compared to WT protein, the mutant SAMHD1 (P275A) did not significantly impair its inhibition of TRAF6-mediated NF-κB activation (Fig. 2F). Taken together, these results suggest that SAMHD1-mediated inhibition of the TRAF6-TAK1 axis contributes to SAMHD1 suppression of the NF-κB signaling.

**Fig. 2.**
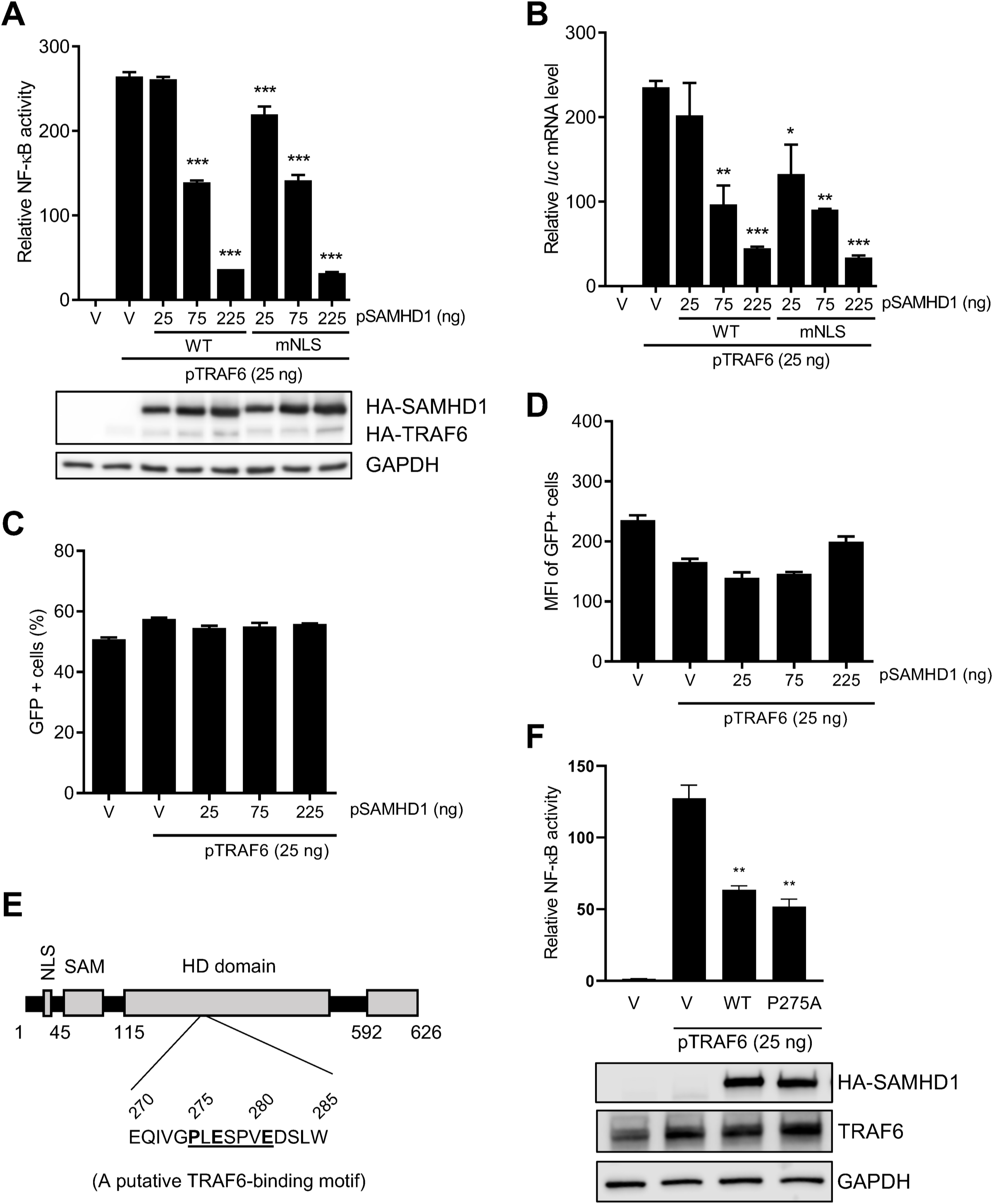
Cytoplasmic SAMHD1 inhibits TRAF6-mediated activation of NF-κB signaling. **(A)** HEK293T cells were co-transfected with the indicated amounts of pRK-HA-TRAF6, wild-type (WT) or mutant NLS (mNLS) pCG-F-HA-SAMHD1 (pSAMHD1), 50 ng pNF-κB-luciferase, and 5 ng of pcDNA3-GFP. The total amount of DNA was maintained through the addition of empty vector (V). Cell lysates were collected for luciferase assay and immunoblotting 24 hr post-transfection. All luciferase values were normalized to 10 µg protein. Relative NF-κB activity was calculated by setting empty vector-transfected cells as 1. HA antibody was used to detect expression of SAMHD1 and TRAF6. GAPDH was used as a loading control. **(B)** Samples from the experiment in (A) were collected to measure *firefly luciferase* (*luc*) mRNA by RT-qPCR. Graphs depict data derived from triplicate samples and error bars represent SEM. Statistical analysis was performed by one-way ANOVA with Dunnett’s multiple comparisons posttest. **(C and D)** Samples from the experiment in (A) were collected to monitor GFP expression by flow cytometry. Graphs depict data derived from triplicate samples and error bars represent SD. Statistical analysis was performed by unpaired t test with Welch’s correction. MFI, mean fluorescence intensity. **(E)** Schematic representation of human SAMHD1 protein highlighting its TRAF6 putative binding motif (^275^PXEXXXE^281^). **(F)** HEK293T cells were co-transfected with pRK-HA-TRAF6 (25 ng), wild-type (WT) or mutant P275A pCG-F-HA-SAMHD1 (pSAMHD1), and 50 ng pNF-κB-luciferase. Empty vector (V) without (the first V) or with pRK-HA-TRAF6 (the second V) were used as negative and positive controls. Cell lysates were collected for luciferase assay (top bar chart) and immunoblotting (bottom blots) at 24 hr post-transfection. An anti-HA antibody was used to detect expression of HA-tagged SAMHD1. GAPDH was used as a loading control. Graphs depict data derived from triplicate samples and error bars represent SD. Statistical analysis was performed by unpaired t test with Welch’s correction. \**P* ≤ 0.05, \*\**P* ≤ 0.01, \*\*\**P* ≤ 0.001.

### TRAF6 KD reduces NF-κB and p38 activation in SAMHD1 KO cells

To further assess the involvement of TRAF6 in SAMHD1-mediated suppression of NF-κB, we utilized control or SAMHD1 KO THP-1 cells to generate vector (LKO) or TRAF6 knockdown (KD) stable cell lines. Immunoblotting confirmed partial KD (50%) of TRAF6 (Fig. 3A). As expected, SAMHD1 KO cells showed higher levels of p-IκBα (Fig. 3A) and *TNF-α* mRNA (Fig. 3B) compared to control cells following IL-1β stimulation. Partial KD of TRAF6 in SAMHD1 KO cells significantly reduced both to the levels observed in control cells (Fig. 3A and 3B), suggesting that TRAF6 is involved in SAMHD1-mediated suppression of canonical NF-κB signaling. The TRAF6-TAK1 axis is central not only to the NF-κB pathway, but also to the c-Jun N-terminal kinase (JNK) and p38 pathways (25, 26). Of note, TLR4 stimulation coordinates signaling which leads to the activation of both the IKK complex and p38 MAPK (35). To determine whether SAMHD1 has the capacity to modulate MAPK activation mediated by upstream signal transduction at the TRAF6-TAK1 axis, we stimulated THP-1 control and SAMHD1 KO cells with LPS for analysis of phosphorylated p38 (p-p38) by immunoblotting. LPS-treated SAMHD1 KO cells showed 1.7-fold enhancement of p-p38 relative to control cells, which was reduced to the level of control (1.9-fold decrease) when SAMHD1 KO was combined with TRAF6 KD (Fig. 3C). Further, elevated *TNF-α* mRNA levels in LPS-stimulated SAMHD1 KO cells were significantly reverted by knocking down TRAF6 (Fig. 3D). These results provide additional evidence that SAMHD1-mediated negative regulation of proinflammatory signaling events involves suppression of the TRAF6-TAK1 axis.

**Fig. 3.**
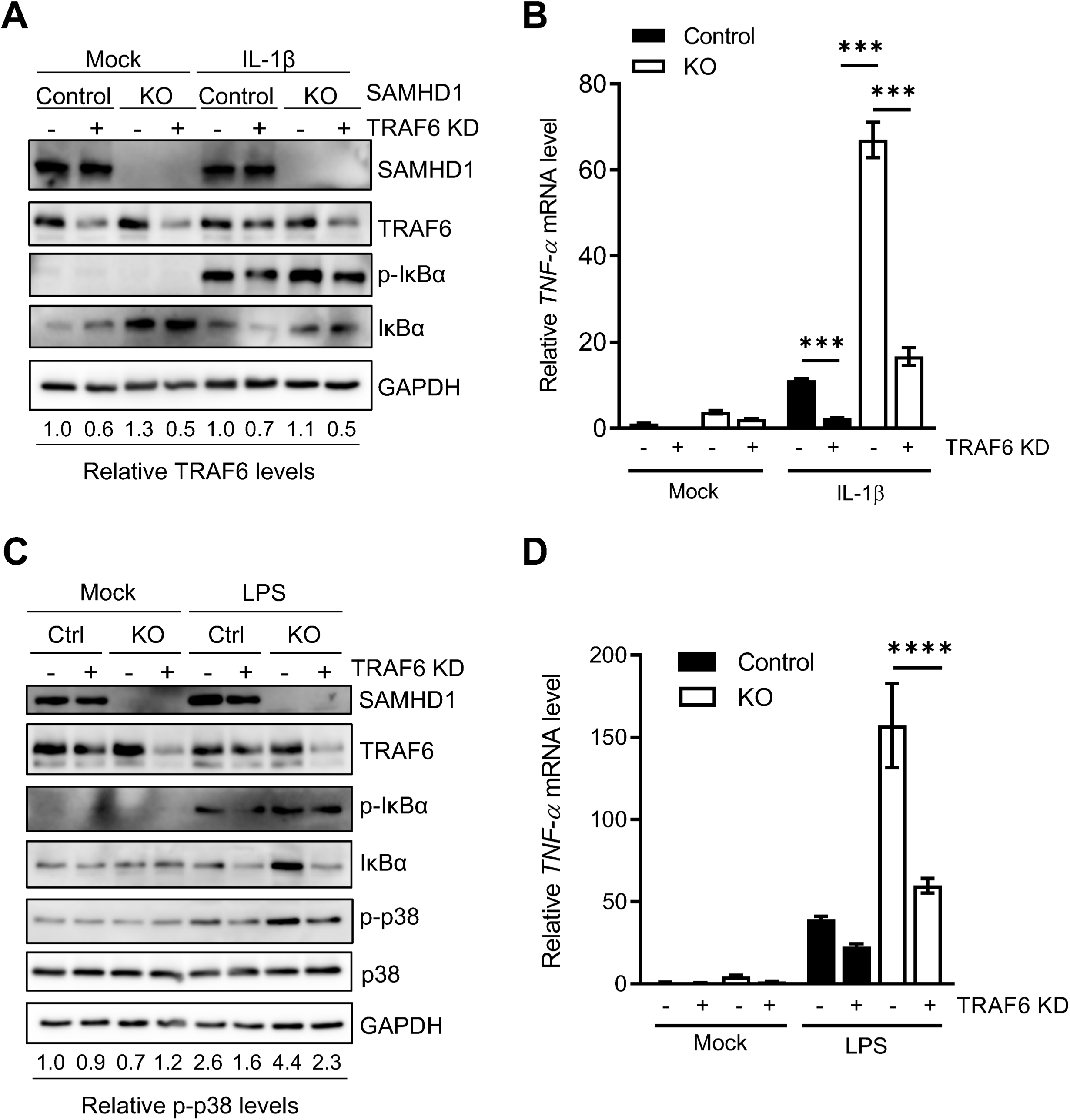
TRAF6 KD reduces NF-κB and p38 activation in SAMHD1 KO cells. **(A)** Stable THP-1 control or SAMHD1 KO cells lines with either control or TRAF6 KD were cultured in standard media and mock-treated or stimulated with IL-1β. Cells were harvested 10 min post-stimulation and lysed for immunoblotting. Relative TRAF6 protein levels were calculated by densitometry analysis. The TRAF6 signal was normalized to GAPDH. Untreated control cells were set as 1. **(B)** Samples for RT-qPCR and mRNA analysis were collected 2 hr post-stimulation. Data represents triplicate samples and error bars depict SEM. Statistical analysis was performed by two-way ANOVA. **(C)** THP-1 control or SAMHD1 KO cells lines with either control or TRAF6 KD were cultured in standard media and mock-treated or stimulated with LPS. After 6 hr, cells were harvested and lysed for immunoblotting. Relative levels of phosphorylated p38 (p-p38) were calculated by densitometry analysis. The p-p38 signal was normalized to total p38 protein and GAPDH. Mock-treated, unstimulated control cells were set as 1. **(D)** *TNF-α* mRNA levels were quantified by RT-qPCR and normalized to spliced GAPDH. The graph depicts data derived from triplicate samples and error bars represent SEM. Statistical analysis was performed by two-way ANOVA with Tukey’s multiple comparisons posttest. \*\*\*\**P* ≤ 0.0001.

### SAMHD1 inhibits NF-kB activation by TAB3/TAK1 or TRAF2

To determine whether SAMHD1 could exclusively inhibit NF-kB activation at the TRAF6 level, we examined the effect of SAMHD1 on NF-kB signaling activated by other intermediate proteins involved in IL-1R/LPS (TAB3/TAK1) and TNF-α-mediated (TRAF2) NF-κB activation. Interestingly, SAMHD1 inhibited NF-kB activation initiated by co-expression of TAB3 and TAK1 proteins in a dose dependent manner (Fig. 4A). SAMHD1 also inhibited TRAF2-mediated NF-κB activation in a dose dependent manner (Fig. 4B). These results suggest that SAMHD1 inhibits NF-kB activation by operating at different levels, which may involve downstream effects at IkBα or IKKε level (17).

**Fig. 4.**
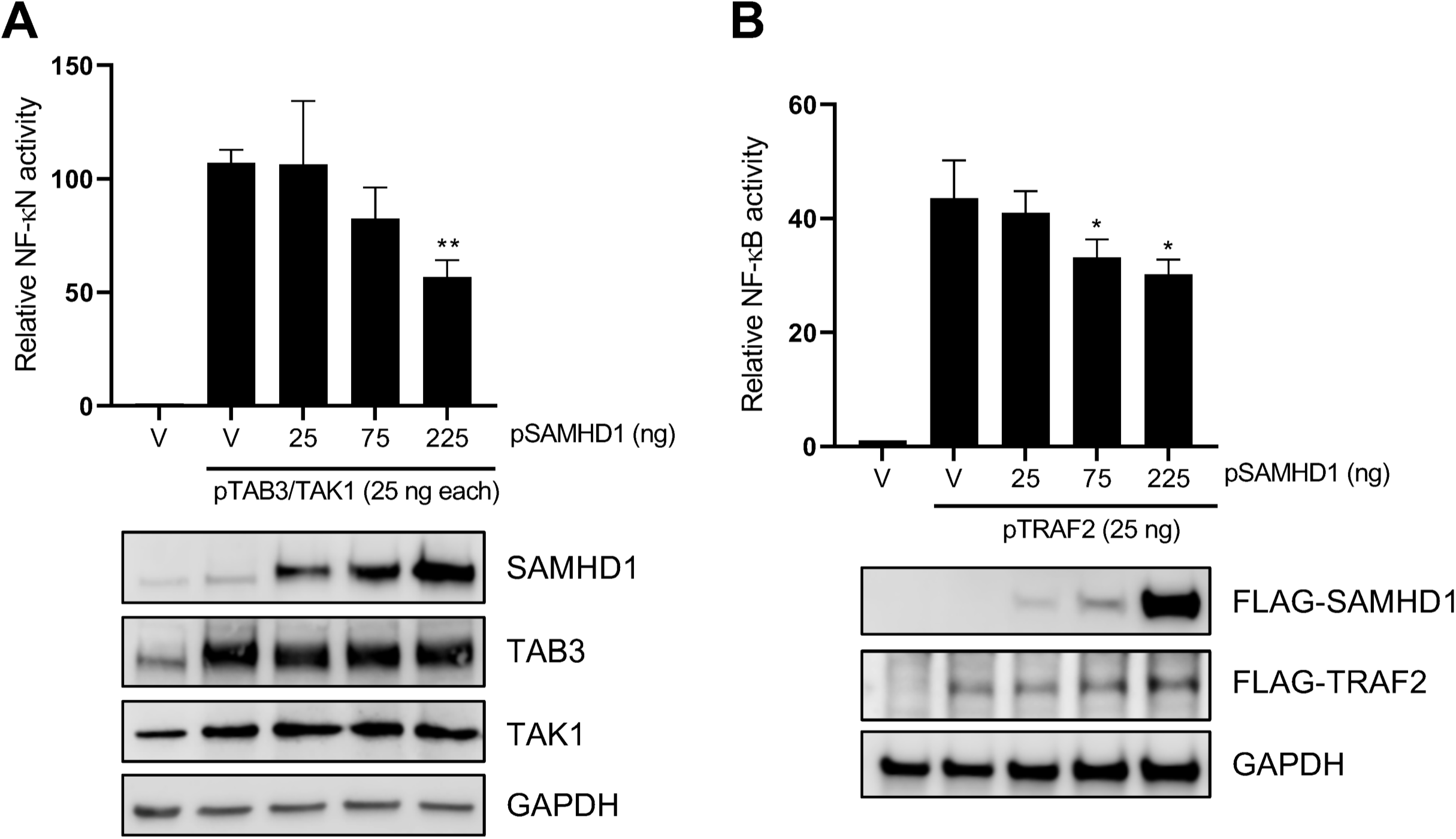
SAMHD1 inhibits NF-κB NF-kB activation by TAB3/TAK1 or TRAF2. **(A)** HEK293T cells were co-transfected with the indicated amounts of pRK-HA-TAB3, pRK-HA-TAK1, wild-type pCG-F-HA-SAMHD1 (pSAMHD1), 50 ng pNF-κB-luciferase and 10 ng of TK-Renilla. Immunoblotting was performed with specific antibodies to the indicated proteins. **(B)** HEK293T cells were co-transfected with the indicated amounts of, pcDNA3.1-F-TRAF2, pCG-F-HA-SAMHD1 (pSAMHD1), 50 ng pNF-κB-luciferase and 10 ng of TK-Renilla. The total amount of DNA was maintained through the addition of empty vector (V). Cell lysates were collected for the luciferase assay and immunoblotting at 24 hr post-transfection. All luciferase values were normalized to 10 µg protein. Relative NF-κB activity was calculated by setting empty vector-transfected cells as 1. Immunoblotting of SAMHD1 and TRAF2 was performed with anti-FLAG antibodies. GAPDH was used as a loading control. Statistical analysis was performed by unpaired t test with Welch’s correction. \**P* ≤ 0.05, \*\**P* ≤ 0.01, \*\*\**P* ≤ 0.001.

### TAK1 inhibition attenuates HIV-1 infection in THP-1 cells lacking SAMHD1 expression

HIV-1 infection does not typically induce potent innate immune responses (36); however, we have reported significant induction of the NF-κB pathway in THP-1 cells lacking SAMHD1 expression (17). As IKK complex activation is a key event in the induction of the NF-κB pathway in response to viral infection, it is not surprising that IKK activity is tightly tuned at multiple levels by regulatory elements, such as the TAK1 protein (37, 38). Therefore, we next evaluated the regulatory impact of SAMHD1 on TAK1 activation during HIV-1 infection. First, THP-1 control and SAMHD1 KO cells were treated with 5Z or Takinib and cells were transduced with VSV-G-pseudotyped single-cycle HIV-1 for 2 hr. Luciferase assay conducted 24 hr post-infection (hpi) showed a 13-fold increase of HIV-1 infection in SAMHD1 KO cells compared to control cells, which was reduced 1.5-fold or 1.8-fold by 5Z or Takinib treatment, respectively (Fig. 5A). Measurement of late reverse transcription (RT) products 6 hr post-transduction showed no difference in copy number between KO and control cells or when cells were treated with either 5Z or Takinib (Fig. 5B). Interestingly, THP-1 cells deficient for SAMHD1 expression showed a 45-fold increase in *luc* mRNA levels compared to control cells, which was reduced by 6.9-fold and 16.7-fold in cells treated with 5Z or Takinib, respectively (Fig. 5C). These data suggest that, in dividing THP-1 SAMHD1 KO cells, enhanced HIV-1 infection is not the result of reverse transcription but rather increased mRNA transcription, and that TAK1 plays a key role in supporting HIV-1 infection. Furthermore, 5Z treatment of HIV-1-transduced SAMHD1 KO cells abrogated the increase in *TNF-α* mRNA transcription (Fig. 5D). Collectively, these data suggest that SAMHD1 may suppress innate immune responses to HIV-1 by diminishing NF-κB activation mediated through TAK1.

**Fig. 5.**
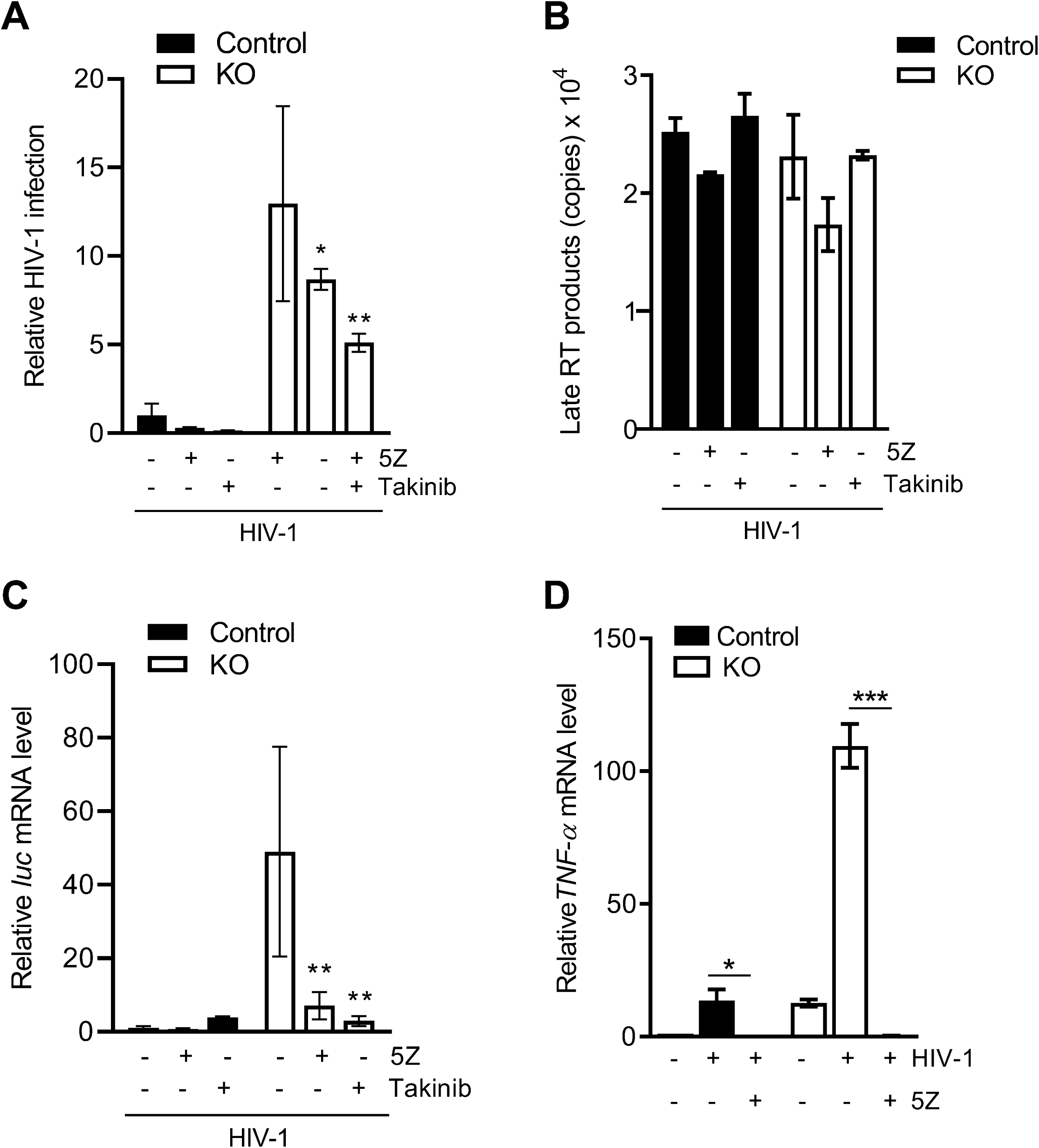
TAK1 inhibition attenuates HIV-1 infection in THP-1 cells lacking SAMHD1 expression. **(A)** THP-1 cells were treated with 5Z (1 µM) for 30 min, Takinib (10 µM) for 2 hr, or DMSO. Inhibitor-containing media was removed prior to 2 hr infection with single-cycle HIV-1-Luc/VSV-G (MOI = 1). Takinib was added and maintained in culture for 24 hr post-infection (hpi). Cells were harvested at 24 hr for luciferase assay. All luciferase values were normalized to 10 µg protein. Relative HIV-1 infection was calculated by setting mock-treated cells as 1. Data represent 4 replicates and error bars show SEM. **(B)** At 6 hpi, late reverse transcription (RT) products were quantified by qPCR assays using samples from the experiment in (A). Serial dilutions 10^8^-10^1^ of a proviral pNL4-3 plasmid were used to calculate late RT products copy numbers. Each biological sample was run in duplicate, and unspliced GAPDH was used for normalization. **(C)** Measurement of mRNA levels was performed from samples in the same experiment described in (A). Cells were harvested at 18 hpi for *luc* mRNA quantification by RT-qPCR. 18S rRNA was used as a normalization control. The graph depicts data derived from triplicate samples with error bars representing SEM. In (A) and (C) statistical analysis was performed by two-way ANOVA with Tukey’s multiple comparisons posttest. **(D)** Cells were treated with 5Z (1 µM) for 30 min or DMSO. Cells were infected for 2 hr with HIV-1-Luc/VSV-G (MOI = 2) in the presence of the inhibitor. *TNF-α* mRNA levels at 2 hpi were quantified by RT-qPCR and normalized to spliced GAPDH. Error bars represent SD of triplicate samples. Statistical significance was calculated by unpaired *t* test. \**P* ≤ 0.05, \*\**P* ≤ 0.01, \*\*\**P* ≤ 0.001.

### TRAF6 contributes to SAMHD1-mediated suppression of HIV-1 mRNA transcription and HIV-1-induced NF-κB activation

We have previously shown that overexpression of SAMHD1 inhibits HIV-1 5’ long terminal repeat (LTR)-driven gene expression (39). It is also known that NF-κB binding to the HIV-1 LTR promoter enhances viral gene transcription (40, 41). We next evaluated whether the increased viral transcription in THP-1 cells lacking SAMHD1 expression was exclusively related to its ability to inhibit the NF-κB pathway. We performed an HIV-1 LTR-driven firefly luciferase (FF-Luc) in HEK293T cells, using WT LTR or a mutant LTR (ΔNF-κB) plasmids, where two NF-κB binding sites have been deleted (42). In agreement with previous reports (43, 44), LTR-driven FF-Luc expression in HEK293T cells was independent of the presence of NF-κB binding sites (Fig. 6A and 6B). Moreover, SAMHD1 overexpression similarly inhibited WT or ΔNF-κB mutant LTR-driven FF-Luc expression in a dose dependent manner (Fig. 6A and 6B), suggesting that SAMHD1 can also suppress HIV-1 LTR-driven gene expression independently of NF-κB binding sites.

**Fig. 6.**
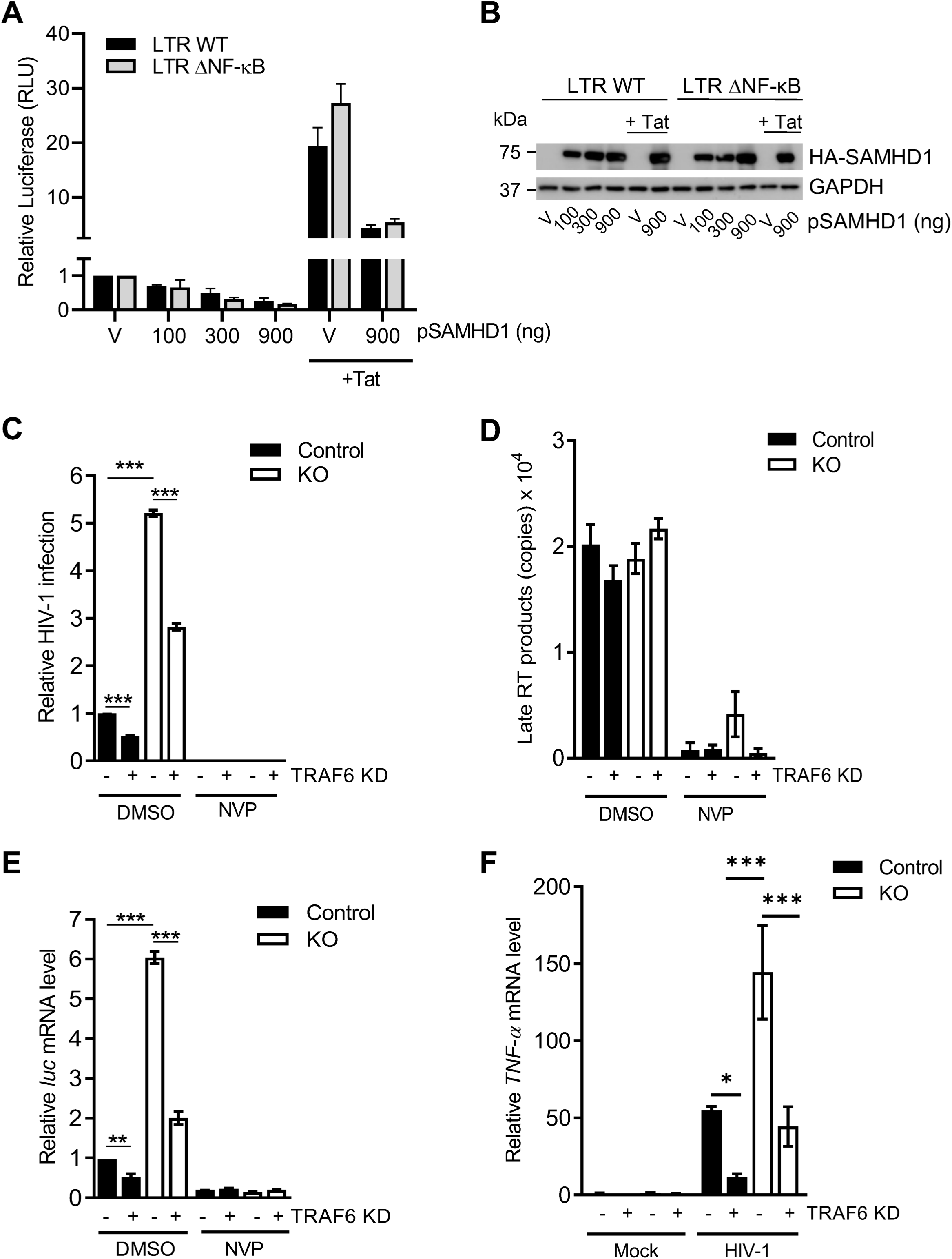
TRAF6 contributes to SAMHD1-mediated suppression of HIV-1 mRNA transcription and HIV-1-induced NF-κB activation. **(A)** HEK293T cells were transfected with an empty vector (V) or increasing amounts of constructs expressing HA-tagged SAMHD1 along with the HIV-1 wild-type (WT) or NF-κB-binding site deleted mutant (ΔNF-κB) LTR-driven FF-Luc reporter. An HIV-1 Tat-expressing plasmid was included to enhance LTR activity via Tat-mediated transactivation. The total amount of DNA was maintained through the addition of empty vector (V). Cell lysates were collected at 24 hr post-transfection for the luciferase assay and immunoblotting. All luciferase values were normalized to 10 µg protein. Relative luciferase expression was calculated by setting empty vector-transfected cells as 1. Statistical analysis was performed by two-way ANOVA. **(B)** Cell lysates from the experiment described in (A) were harvested at 24 hr post-transfection for immunoblotting. An anti-HA antibody was used to detect expression of HA-tagged SAMHD1. GAPDH was used as a loading control. **(C)** THP-1 control or SAMHD1 KO cells in the nevirapine (NVP) group were pre-treated with NVP (5 µM) for 1-2 hr prior to 2 hr transduction with single-cycle HIV-1-Luc/VSV-G (MOI = 1). NVP was maintained in the medium throughout the infection and subsequent culture. Cells were harvested at 24 hpi for luciferase assay. All luciferase values were normalized to 10 µg protein. Relative HIV-1 infection was calculated by setting control cells without TRAF6 KD as 1. **(D)** At 6 hpi, late RT products were quantified by qPCR assays using samples from the experiment in (A). Serial dilutions 10^8^-10^1^ of an HIV-1 proviral pNL4-3 plasmid were used to calculate late RT copy numbers. Each biological sample was run in duplicate, and unspliced GAPDH was used for normalization. The data in (A-C) depict triplicate samples and error bars represent SEM. **(E)** Measurement of mRNA levels was performed from samples in the same experiment described in (A). Cells were harvested at 18 hpi for *luc* mRNA quantification by RT-qPCR. Spliced GAPDH was used as a normalization control. **(F)** Cells were transduced with HIV-1-Luc/VSV-G (MOI = 1) for 2 hr, then further cultured prior to collection at 2 hpi for detection of *TNF-α* mRNA by RT-qPCR. Spliced GAPDH was used as a normalization control. Statistical analysis was performed by two-way ANOVA. \**P* ≤ 0.05, \*\**P* ≤ 0.01, \*\*\**P* ≤ 0.001.

Our results showed that in dividing THP-1 cells, SAMHD1 suppress viral transcription by diminishing TAK1-mediated NF-κB activation (Fig. 5). To further examine the role of the SAMHD1 in HIV-1 transcriptional inhibition mediated by TRAF6-TAK1 axis, the effect of TRAF6 KD on HIV-1 infection and viral mRNA transcription in THP-1 cells was assessed. Indeed, TRAF6 KD reduced HIV-1 infection at 24 hpi by ∼2 fold in both SAMHD1 KO and control cells (Fig. 6C). HIV-1 late RT products were similar in KO and control cells, irrespective of TRAF6 KD and they could not be detected in nevirapine (NVP)-treated cell samples (Fig. 6D). In contrast, TRAF6 KD reduced viral *luc* mRNA by 3-fold in SAMHD1 KO and by 2-fold in control cells (Fig. 6E), suggesting that TRAF6 is an important cellular component for HIV-1 infection at a replication stage between late RT and mRNA transcription. We further evaluated whether TRAF6 is involved in NF-κB activation induced by HIV-1. As expected, SAMHD1 KO cells had 2.6-fold higher levels of *TNF-α* mRNA compared to control cells (Fig. 6F). Of interest, TRAF6 KD was sufficient to reduce *TNF-α* mRNA levels in SAMHD1 KO cells to levels comparable with control cells. Altogether, these results suggest that SAMHD1 contributes to suppressing the innate immune responses initiated through the TRAF6-TAK1 signaling complex during single-cycle HIV-1 infection.

## Discussion

The antiviral innate immune response is regulated at multiple steps in the signaling cascade (45-47). We have previously revealed a novel role for a well-described HIV-1 restriction factor, SAMHD1, as a negative modulator of the IFN-I and NF-κB pathways (17). The NF-κB pathway is activated through the upstream kinase TAK1. We found that SAMHD1 expression is sufficient to reduce TAK1 activation and that pharmacological inhibition of TAK1 reverts SAMHD1-mediated suppression of NF-κB signaling. Furthermore, SAMHD1 can reduce NF-κB activation initiated by TRAF6 overexpression, which is independent of receptor stimulation. This observation may have important implications because the TRAF6-TAK1 axis is a central point from which several signaling pathways branch off, including the NF-κB, MAPK, phosphoinositide 3-kinase (PI3K), and interferon regulatory factor (IRF) pathways (48, 49). Indeed, SAMHD1 has been implicated in the PI3K/AKT/IRF3 signaling (50), hinting at a possible global mechanism by which SAMHD1 protein modulates key signaling events. In addition, our results showed that SAMHD1 is able to reduce MAPK p38 phosphorylation, providing further evidence of a broader role for SAMHD1 in modulating several signaling cascades that could affect cellular transcription, inflammation, and innate immune activation in response to HIV-1 infection and proinflammatory stimuli (Fig. 7).

**Fig. 7.**
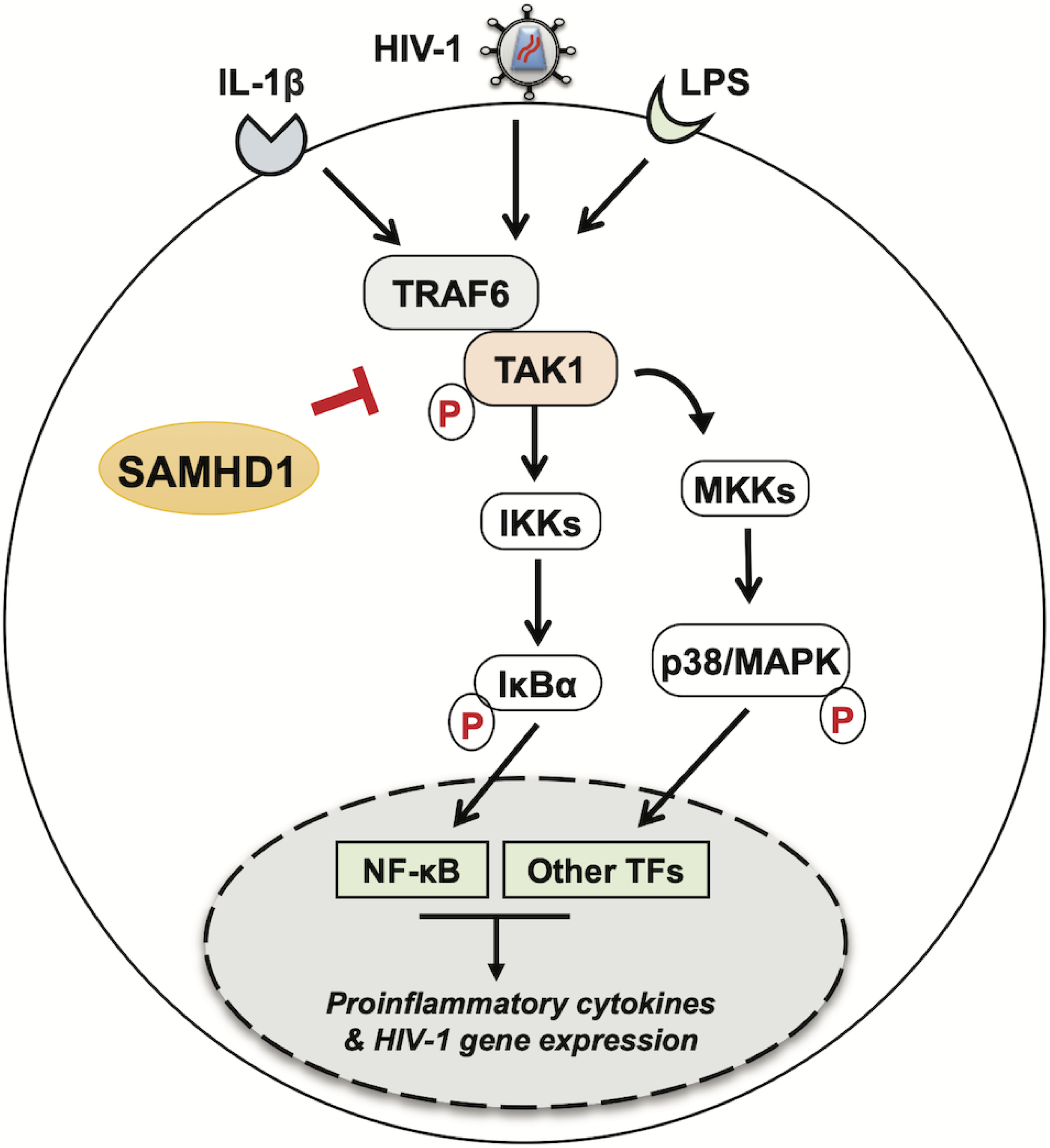
SAMHD1 inhibits signal transduction mediated by the TRAF6-TAK1 complex during HIV-1 infection or cytokine stimulation. Cytoplasmic SAMHD1 suppresses the NF-κB and p38 MAPK pathways, which are activated through signaling cascades that branch from the TRAF6-TAK1 axis. SAMHD1 inhibits phosphorylation of the IκBα and p38 proteins induced by cytokine stimulation (IL-1β) or recognition of pathogen-associated molecular patterns (LPS) by cells, as well as NF-κB-mediated gene expression by reducing *TNF-α* mRNA synthesis. SAMHD1 suppresses innate immune responses to HIV-1 infection through negative modulation of TRAF6-TAK1 signaling, as SAMHD1-deficient cells infected with HIV-1 show enhanced production of viral mRNA transcripts, which is reduced by TRAF6 silencing or pharmacological inhibition of TAK1. For simplicity, only cytoplasmic SAMHD1 is shown. The letter p in red indicates phosphorylation.

SAMHD1 is most well-known for restricting HIV-1 infection in non-dividing cells where its dNTPase activity limits the dNTP levels below the threshold required for efficient HIV-1 late reverse transcription (22, 51, 52). We previously reported that SAMHD1 KO in dividing THP-1 cells increases the dNTP pool and HIV-1 infection (53). However, THP-1 cells are not restrictive to HIV-1 infection and it is still unclear whether SAMHD1-mediated HIV-1 restriction can be solely attributed to its dNTPase function (54). In this study we showed that SAMHD1 additionally limits NF-κB-dependent HIV-1 transcription in dividing THP-1 cells through the TRAF6-TAK1 axis. Further studies are required to delineate the precise molecular mechanisms through which SAMHD1 exerts this effect. Interestingly, HIV-1 viral proteins have been found to enhance viral replication through manipulation of NF-κB signaling. HIV-1 Vpr activates TAK1 and hence NF-κB to enhance LTR activation (55) and HIV-1 envelope glycoprotein gp41 interacts with TAK1 and induces NF-κB activation to facilitate viral replication (56). Our studies were carried out in the absence of these viral components, which raises the possibility that viral proteins may also counteract SAMHD1-suppression of NF-κB in order to maintain HIV-1 replication. Furthermore, TRAF6 has been shown to regulate HIV-1 production in macrophages (57) and knockout of microRNA-146a, which targets TRAF6, reduces HIV-1 infection and reactivation of latently infected cells (58), providing further evidence that TRAF6 plays an important role in HIV-1 infection.

Our current and previous results demonstrated that nuclear localization of SAMHD1 is dispensable for suppression of NF-κB activation in HEK293T cells and differentiated THP-1 or U937 monocytic cell lines (20). These observations support the notion that a cytoplasmic portion of SAMHD1 may specifically function to block proteins involved in NF-kB signaling, such as TRAF6/2, TAB3, and/or TAK1. While the SAMHD1 protein contains a predicted TRAF6 binding domain, we found that disruption of this domain did not impair inhibition of TRAF6-mediated NF-κB activation by SAMHD1. Our immunoprecipitation studies indicated that SAMHD1 does not interact with TAK1 and/or TRAF6 in HEK293T or THP-1 cells (data not shown), suggesting that the suppressive effect of SAMHD1 does not require its interaction with these cellular proteins.

Together, the results from this study provide new insights into the molecular mechanisms used by SAMHD1 to reduce host cell immune responses during HIV-1 infection or proinflammatory stimulation, thus highlighting potential therapeutic approaches for the control of HIV-1 infection and inflammatory diseases.

## Materials and Methods

### Plasmids

The plasmid encoding HA-tagged RK-TRAF6 was a kind gift from Dr. S. Chen. The plasmids encoding for TAB3 and TRAF2 were a kind gift from Dr. C. Maluquer de Motes. The plasmids encoding for WT-LTR FFLuc and ΔNF-κB LTR-FFLuc were provided by Dr. J. Wang and were described in (42). Plasmid encoding shRNA targeting human TRAF6 in the pLKO.1 lentiviral vector was purchased from Dharmacon (RHS3979-201739625). The pLKO.1 vector control and the pNF-κB-luciferase vector (PRDII4–luc in the pGL3 vector) were previously described (17). The pcDNA3-GFP vector was previously described (59). The pCG-FLAG-HA plasmids (60) encoding the SAMHD1 mutant NLS (mNLS) and the P275A mutants were generated using a QuikChange Site-Directed Mutagenesis Kit (Agilent Technologies) according to the manufacturer’s protocol using the following primers: 5’-GAG CAG CCC TCC GCG GCT CCC GCT TGC GAT GAC AGC (mNLS) (20) and 5’-GGA ACA AAT TGT AGG AGC ACT TGA ATC ACC TGT C (SAMHD1 P275A). The pCG-FLAG-HA-SAMHD1 plasmid was a kind gift from Dr. J. Skowronski. Single-cycle, VSV-G-pseudotyped luciferase reporter HIV-1 (pNL4-3E^-^R^+^) was a kind gift of Dr. N. Landau. psPAX2 and CMV promoter-driven VSV-G (pMD2.G) were gifted from Dr. P. Spearman.

### Cell culture

HEK293T cells were purchased from the American Type Culture Collection (ATCC) (60). THP-1 control or KO cells were described previously (53). THP-1 control or SAMHD1 KO cells stably expressing empty vector or TRAF6 shRNA were generated by spinoculation with concentrated lentiviral vectors followed by 1 μg/mL puromycin selection. Transduced THP-1 cell lines were cultured in RPMI 1640 (ATCC) supplemented with 10% fetal bovine serum, 100 U/mL penicillin, 100 μg/mL streptomycin, and 1 μg/mL puromycin. HEK293T cells were cultured in DMEM with 10% fetal bovine serum, 100 U/mL penicillin and 100 μg/mL streptomycin. All cell lines were maintained at 37 °C, 5% CO_2_ and confirmed free from mycoplasma contamination using the universal mycoplasma detection kit (ATCC 30-101-2K).

### IL-1β treatment, (5Z)-7-Oxozeaenol and Takinib-mediated inhibition of TAK1 in THP-1 control or SAMHD1 KO cells

Dividing THP-1 cells were treated with 10 ng/mL recombinant human IL-1β (Peprotech) for the times indicated. Mock-treated cells were treated with media only. After the indicated treatment time, cells were collected for either RNA extraction and qPCR analysis or cell lysis and immunoblotting. (5Z)-7-Oxozeaenol (5Z) was purchased from Sigma-Aldrich (O9890). Stock solutions of inhibitor were stored in DMSO. A final concentration of 1 µM was used to pre-treat cells for 30 min prior to stimulation and downstream processing. Cells in the untreated group were given an equal volume of DMSO. Takinib (Sigma-Aldrich - SML2216) was also stored in DMSO. A 10 µM final concentration was used for treatment of cells. 2 hr pre-treatment was performed prior to downstream processing. Longer treatment periods are indicated in the figure legends where appropriate. For detection of phosphorylated TAK1 (p-TAK1), cells were grown in low glucose (5.5 mM) media for 48 hr prior to additional experimental treatments.

### LPS treatment of cells

THP-1 cells were treated with 100 ng/mL LPS from *Escherichia coli* (Sigma-Aldrich – L6895) or media only (mock treatment) for 6 hr prior to collection for RNA extraction and RT-qPCR or cell lysis and immunoblotting (17).

### Antibodies and immunoblotting

Cells were lysed in cell lysis buffer (Cell Signaling) containing protease inhibitor cocktail (Sigma-Aldrich) and phosphatase inhibitor cocktail 3 (Sigma-Aldrich), 10 mM NaF and 1 mM PMSF. Cell lysates were analyzed by a BCA assay (Pierce) for protein quantification, and equal amounts of protein were loaded for SDS-PAGE. Immunoblotting was performed as described using the following specific antibodies: HA (Biolegend, #901501), IκBα (Abnova, #MAB0057), p-IκBα and TRAF6 (Cell Signaling, #9246S, #8028), p-p38 and p38 MAPK (Cell Signaling, #4511T, #8690T), GAPDH (Bio-Rad, #AHP1628), SAMHD1 (Abcam, #67820 or ProSci, #8007), TAB3 (#14241S), TAK1 (#5206S) and pTAK1 (T187) (#4536S) and Flag (Sigma # F1804). Detection of GAPDH protein was used as a loading control. For densitometry, low exposure images were analyzed using ImageJ or Image Studio Lite, and the signal for the target protein was normalized to that of GAPDH (17).

### HIV-1 infection

HIV-1 infection of stable THP-1 control and KO cells lines was performed using single-cycle NL4-3E^-^R^+^ luciferase virus pseudotyped with VSV-G. Virus stocks were treated with 40 U/mL DNase I (Invitrogen) for 1 hr at 37°C before they were used to infect cells for 2 hr at an MOI of 0.5-2, or as indicated. Pre-treatment of cells with 5 µM NVP (NIH AIDS Reagent Program, #4666) was performed where described. After 2 hr infection, media containing virus was removed and cells were re-plated in fresh media with or without inhibitors, as described in the figure legends, for the indicated times.

### RNA extraction and real-time quantitative RT-PCR assay

Cells were collected and total RNA was extracted using the RNeasy Mini Kit (Qiagen) or Aurum™ Total RNA Mini Kit (Bio-Rad). RNA concentrations were determined by Nanodrop and equal amounts of RNA were reverse transcribed into cDNA using the First-Strand Synthesis Kit IV (Invitrogen) or iScript cDNA Synthesis Kit (Bio-Rad). Equal volumes of cDNA were used for iTaq Universal SYBR Green (Bio-Rad)-based qPCR detection. Spliced *GAPDH* or *18S rRNA* was used as normalization controls, as indicated in the figure legends, and relative mRNA levels were calculated using the 2^-ΔΔCT^ method. Primer sequences for *TNF-α* and *firefly luciferase* target genes were previously reported (17, 32).

### DNA extraction and real-time quantitative PCR of HIV-1 late reverse transcription products

Cells were collected and total genomic DNA was extracted using the DNeasy Blood and Tissue Kit (Qiagen). DNA concentrations were determined by Nanodrop and equal amounts were used for iTaq Universal SYBR Green (Bio-Rad)-based qPCR detection. Copy number was determined using a proviral DNA plasmid (pNL4-3) as a standard. Unspliced GAPDH was used for normalization. Primer sequences for late reverse transcripts have been described previously (17, 32).

### NF-κB luciferase assay in HEK293T cells

HEK293T cells were co-transfected by polyethylenimine (PEI) with pRK-TRAF6 and WT or mNLS SAMHD1. Empty vector controls were used to maintain equal amounts of transfected DNA. A GFP construct (pcDNA3-GFP) was also transfected to monitor transfection efficiency. At 24 h post-transfection, cells were collected for luciferase assay using the Luciferase Assay System (Promega) according to the manufacturer’s protocol (17). The BCA assay was also performed to normalize luciferase results. Separate wells were collected and lysed for immunoblotting analysis.

### Statistical analysis

Statistical analysis was performed as detailed in each figure legend. Data were analyzed using the GraphPad Prism software (Version 5 or 8). Statistical significance was defined as *P* ≤ 0.05.

## Acknowledgements

We thank members of the Wu laboratory and Dr. J. Yount for helpful discussions; Drs. S. Chen, C.M. de Motes, N. Landau, J. Skowronski, and P. Spearman for reagents; H. Wang from the Ohio State University (OSU) Comprehensive Cancer Center Genomic Core Facility for processing microarray samples and Dr. L. Yu from the OSU Center for Biostatistics for Ingenuity Pathway Analysis. This work was supported by NIH Grants R01AI141495 and R01AI150343 to L.W. V.M was supported by C. Glenn Barber Funds from the OSU College of Veterinary Medicine. Nevirapine was obtained through the NIH AIDS Reagent Program.

